# Expanded Scope of Practice Fellowships for Radiologists: a Survey of Interest Amongst Current Canadian Radiology Residents

**DOI:** 10.1101/626770

**Authors:** Joseph J Barfett, Errol Colak, Christopher U Smith, Paraskevi Vlachou, Aren Mnatzakanian, Hui Ming Lin, Eric Bartlett, Alan Moody

**Author notes:** **Corresponding Author:** Joseph John Barfett, MD, MSc Department of Medical Imaging, St. Michael’s Hospital 30 Bond St., Toronto, Ontario, M5B 1W8. This research did not receive any specific grant from funding agencies in the public, commercial, or not-for-profit sectors.

## Abstract

**Rationale and Objectives:** Radiology residents acquire a diverse educational experience and skill set, including a general internship year, which may enable the direct management of patients. In order for radiology residents to define new scopes of practice, however, additional fellowship training may in certain instances be warranted.

**Materials and Methods:** Using the Canadian family medicine Enhanced Skills Program as a model, we conducted a Canada-wide survey of radiology residents to assess interest in additional fellowship training to expand their scope of practice.

**Results:** Our results indicate that a majority of residents (69.2%) would like to routinely see patients in clinic and more than half (52%) are willing to undergo an additional year of fellowship to enhance their skill set. The most popular choices for such fellowships were sports medicine (22.8%), emergency medicine (19.6%) and vascular medicine (18.5%). In addition, a majority (52.9%) of residents felt capable of offering incidentaloma clinics without additional training beyond their core radiology residency.

**Conclusion:** Traditional diagnostic and interventional radiology fellowships must be reconsidered to reflect the interests and capabilities of modern radiology trainees. Expansion of training options into the domain of direct patient management will likely prove popular among current residents.

## Introduction

Radiologists do not disdain working directly with patients and senior medical students do not list isolation from patients as a factor influencing their decision to enter radiology residencies [1]. Indeed the lack of patient interaction and loss of control over patient referrals may be counterproductive to our specialty and one of the reasons interest in radiology residency is declining amongst medical students [2–4]. Although the rise of interventional radiology as a distinct specialty [5] has partially filled this void, there remains significant opportunity to expand the future scopes of practice available to motivated radiology trainees. Medicine, medical imaging and especially medical education and radiology residency training are constantly evolving and with change comes both risk and opportunity. Family medicine programs have responded particularly well to such challenges and currently offer numerous fellowships to extend the scope of practice of their trainees [6]. Residency administrators should take seriously the reality that radiology residents aspire to direct patient management roles in the health system and design training programs appropriately.

We undertook a national survey of radiology residents in Canada to determine their interest in completing further fellowship training in expanded scope of practice fellowships (referred to herein as “clinical fellowship”). We define a “clinical fellowship” as distinct from traditional diagnostic imaging and interventional fellowships and include low risk anesthesia, non-surgical breast disease, cancer screening, chronic pain, clinician scholar, diabetes and wound care, emergency medicine, hospital medicine, incidentaloma management, low risk obstetrics, medical oncology, palliative care, public or environmental health, renal stone disease, sports and exercise medicine, vascular/atherosclerosis medicine and “other”, where we queried resident’s interest. These fellowships are indeed based on the Enhanced Skills Program offered to family physicians in Canada [6–8] where training time varies between individual programs but does not exceed one year in addition to the regular 2-year family medicine program.

The management of incidentalomas is a subject of particular relevance. It is not necessarily the case that radiologists require additional training to offer incidentaloma clinics which include minimally invasive biopsy and management services. An incidentaloma specialization enables broad access to patient management. Interest amongst trainees in incidentalomas and their management and the need for further training are specifically examined in our survey.

We concluded the survey by requesting resident input into the issue of further clinical training. We present the anonymized comments to fuel further discussion on this subject.

## Methods

With research ethics board approval, an electronic survey was distributed to the 13 English speaking radiology residency program directors in Canada with a request to distribute the link to the residents within their program. Approximately 325 resident physicians over the 5 post-graduate years were surveyed as per the Canadian Resident Matching Service (CaRMs) website [8], not taking into account program transfers or alternative funding mechanisms.

### Survey questions are presented below

1. What is your gender? (options: female, male, other)
2. What is your age? (options: <25, 25-29, 30-34, 35+)
3. Where did you receive your medical degree? (options: all Canadian schools were listed in addition to an outside of Canada option)
4. What Canadian diagnostic radiology program are you currently enrolled in? (options: all Canadian programs were listed)
5. What is your current Post Graduate Year (PGY) level? (options: PGY1, 2, 3, 4, 5 and other)
6. Was diagnostic radiology your first choice in the Canadian Residency Matching Service match (CaRMS)? (options: yes or no)
7. Do you plan to pursue a diagnostic radiology or interventional radiology fellowship training after residency?(options: yes, no, uncertain)
8. What would be your preferred diagnostic radiology or interventional radiology fellowship? (options: unsure, abdominal/body, breast, cardiothoracic, cross-sectional, emergency and trauma, head and neck, interventional, Musculoskeletal (MSK), neuro-diagnostic, neuro-interventional, nuclear medicine, obstetrical imaging, pediatric, women’s imaging, other) Family Medicine residents have form many years completed additional fellowships following their two-year residency to support more focused clinical practice (ER, palliative care, pain medicine, etc). The following questions refer specifically to these types of clinical fellowships which are focused on direct patient interaction and management. For the purposes of this survey, these are distinct from conventional diagnostic imaging and interventional fellowships and we refer to them as “clinical fellowships”.
9. Would you be interested in completing a formal “clinical fellowship”? (scale between 0-100%)
10. Would you prefer to complete a clinical fellowship: (options: before, after or instead of a diagnostic or interventional radiology fellowship)
11. Assuming a one-year time commitment, please rank your top three choices for “clinical fellowship”: Please rank up to 3 choices, with the 1st choice being your most preferred. If you would be interested in fellowships which are not listed please include in the “other” section, in order of preference. (options: fellowships listed as per table 1)
12. Would you be interested in seeing patients in clinic as a staff radiologist? (options: yes, no, unsure)
13. In ideal circumstances, what percentage of your work hours would you like to dedicate to clinic? (scale between 0-100%)
14. Do you think it is appropriate for radiologists to offer “incidentaloma” clinics as a means of supporting clinicians who would prefer specialist follow-up and management of incidental findings on imaging tests? (options: yes, no, unsure)
15. Would you be comfortable offering an incidentaloma clinic after residency, without further fellowship training, if you were supported by ACR guidelines? (options: yes, no, unsure)
16. Do you think practicing radiologists with or without conventional fellowship training should be permitted to access clinical fellowships (as we have defined them in this survey) to expand their scope of practice? (options: yes, no, unsure)
17. Thank you for completing this survey. Feel free to provide any additional comments.

**Table 1:** Choice of “clinical fellowship” to be pursued in addition to or in lieu of a traditional diagnostic or interventional radiology fellowship. The most popular choices include emergency medicine, sports and exercise medicine and vascular/atherosclerotic medicine.

| “Clinical Fellowship” Choice if an Additional Year of Training were Undertaken | Number Selecting as First Choice | Number Selecting as Second Choice | Number Selecting as Third Choice | Total Selecting each Fellowship |
| --- | --- | --- | --- | --- |
| Anesthesia | 3 | 3 | 4 | 10 |
| Breast Diseases (non-surgical or benign) | 5 | 3 | 3 | 11 |
| Cancer screening | 3 | 7 | 10 | 20 |
| Chronic pain | 5 | 7 | 3 | 15 |
| Clinician Scholar | 3 | 0 | 5 | 8 |
| Diabetes / wound care | 0 | 0 | 1 | 1 |
| Emergency Medicine | 18 | 15 | 8 | 41 |
| Hospital Medicine | 1 | 2 | 3 | 6 |
| Incidentaloma management | 3 | 8 | 10 | 21 |
| Low risk obstetrics | 1 | 2 | 1 | 4 |
| Medical oncology | 8 | 9 | 8 | 25 |
| Palliative Care | 0 | 3 | 3 | 6 |
| Public or environmental health | 2 | 5 | 3 | 10 |
| Renal stone disease | 0 | 2 | 1 | 3 |
| Sports and exercise medicine | 21 | 17 | 9 | 47 |
| Vascular / atherosclerotic medicine | 17 | 7 | 4 | 28 |
| Other | 2 | 0 | 2 | 4 |

## Results

Response rates for the survey questions were as follows: Question 1-115 responses received, 2-115 responses received, 3-113 responses received, 4-110 responses received, 5-110 responses received, 6-96 responses received, 7-94 responses received, 8-94 responses received, 9-104 responses received, 10-95 responses received, 11-104 responses received, 12-104 responses received, 13-104 responses received, 14-16 responses received.

Where residents indicated their program of origin, responses were received from 11/13 programs queried. In total, 115 residents responded, 14 from eastern Canada, 10 from Quebec, 49 from Ontario, 8 from the prairies and 33 from western Canada. The largest response was from the University of Toronto (29.2%). Two residents did not identify their residency program of origin. Of these residents, 14 completed medical education at schools outside Canada (12.2%) and 99 were Canadian trained (77.8%). Two residents did not identify where they completed medical school.

79.1% of survey respondents were male. The distribution of ages is shown in figure 1a. The distribution of training years is shown in figure 1b. 92.0% of respondents chose diagnostic radiology as their first choice in the match. 90.0% of respondents planned on pursuing traditional diagnostic or interventional radiology fellowship training, 2.7% did not plan to pursue a fellowship and 7.3% were uncertain. The distribution of diagnostic or interventional radiology fellowships of interest to the group is shown in figure 2.

**Figure 1:**
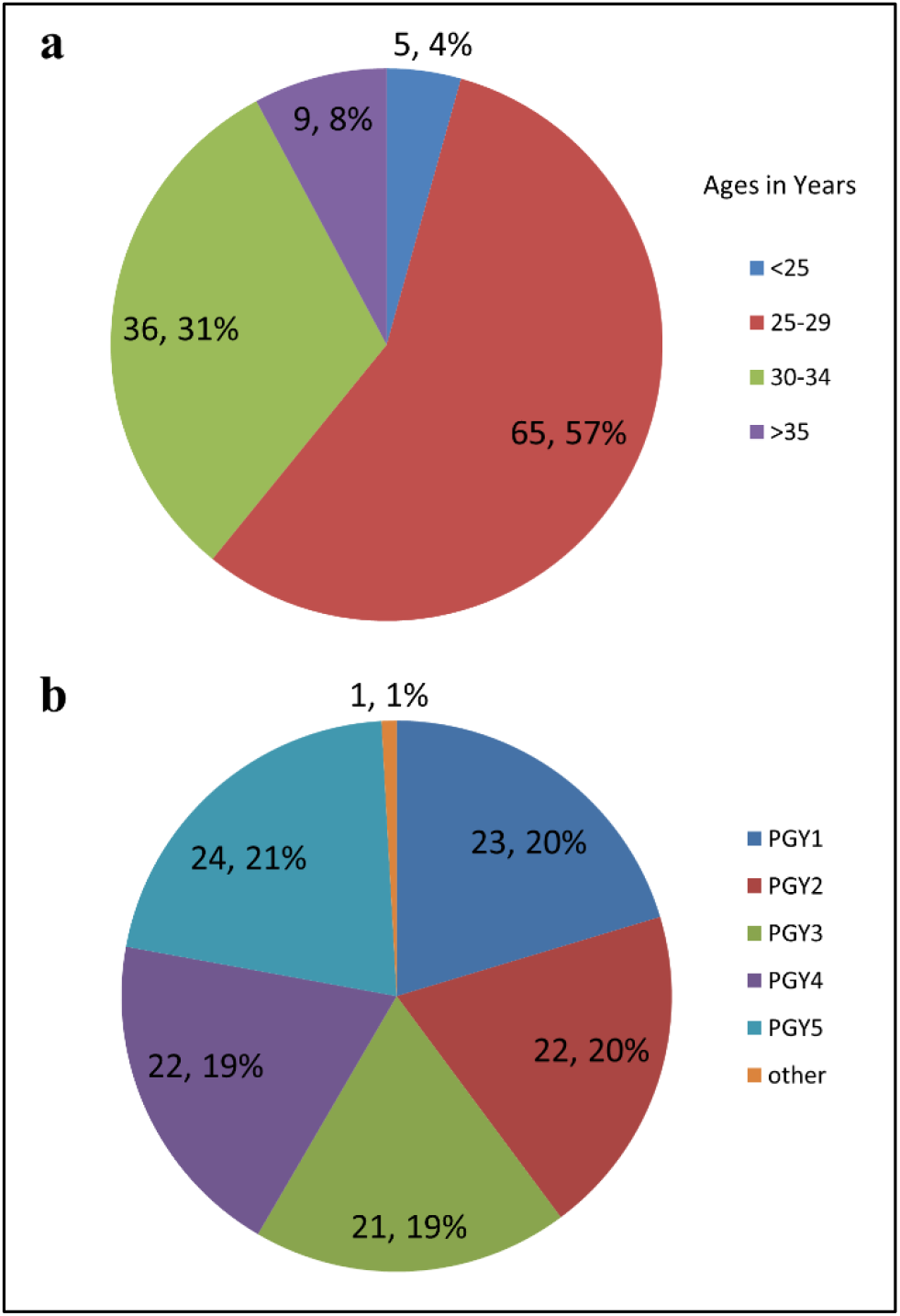
Ages (a) and year level (b) of survey respondents. PGY-post graduate year

**Figure 2:**
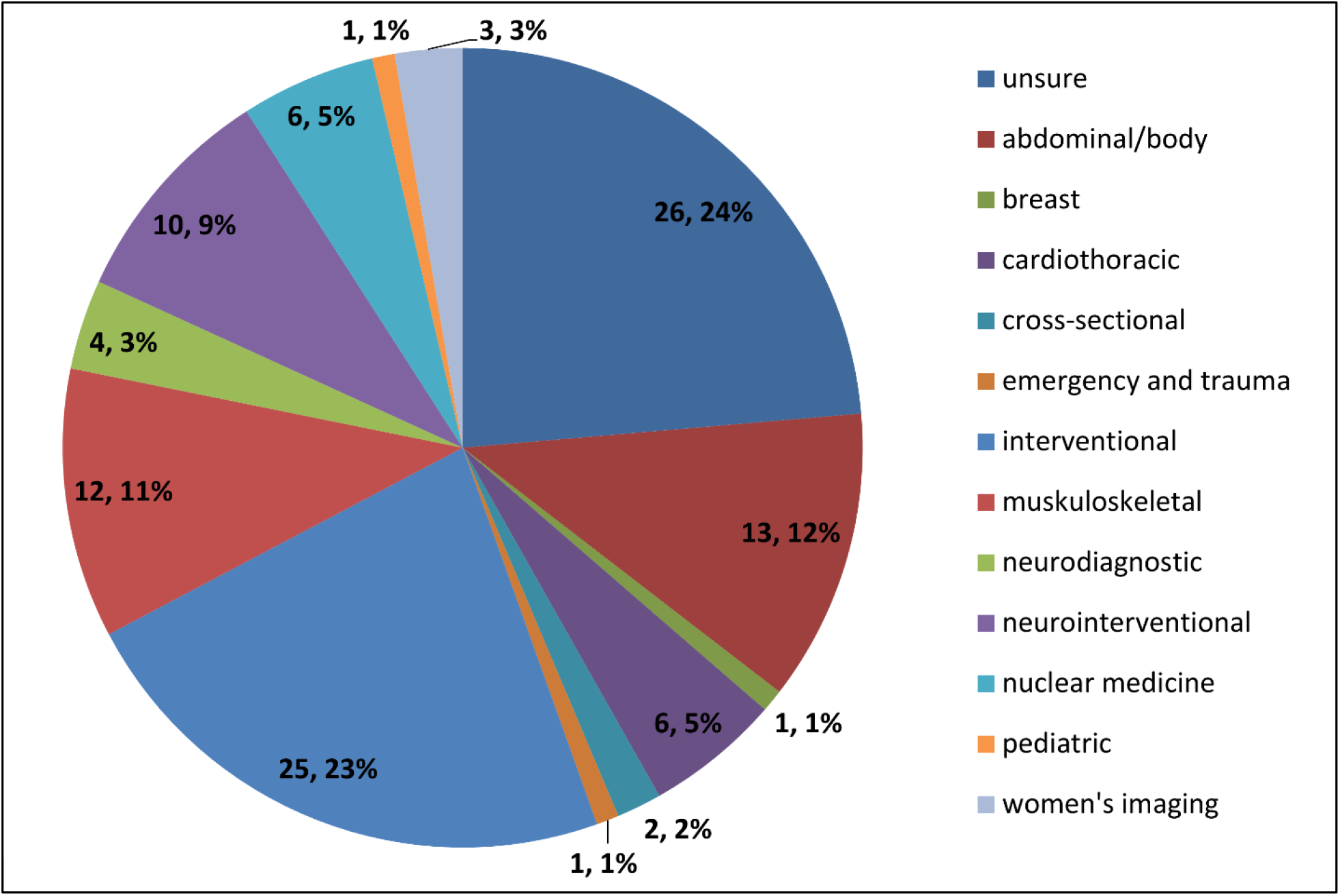
Traditional fellowship choices of the respondents opting for conventional fellowship training

69.2% of respondents indicated that they would like to see patients in clinic as a component of their practice. Respondents indicated they would choose to dedicate a mean of 23.8% (+/− 14.8%) of their work week to clinic. 10 respondents (10.5%) would opt to spend 2 or more days per week seeing patients in clinic and 35 (36.8%) between 1 and 2 days seeing patients in clinic.

When asked to rate their interest in pursuing a “clinical fellowship” on a scale of 1-100, 52/104 expressed >50% interest (50%), 46 expressed >60% interest (44.2%), 33 expressed >70% interest (31.7%), 20 expressed > 80% interest (19.2%), 16 expressed > 90% interest (15.4%) and 14 expressed 100% interest (13.5%). Of those interested in clinical fellowship, 63.8% would opt to complete the training after their conventional diagnostic or interventional fellowship, 21.3% before their diagnostic or interventional fellowship and 14.9% would opt for a clinical fellowship instead of a diagnostic or interventional fellowship. Choices for “clinical fellowship” are shown in table 1 and figure 3 with the most popular choices for such fellowships including sports medicine (22.8% of respondents), emergency medicine (19.6%) and vascular medicine (18.5%).

**Figure 3:**
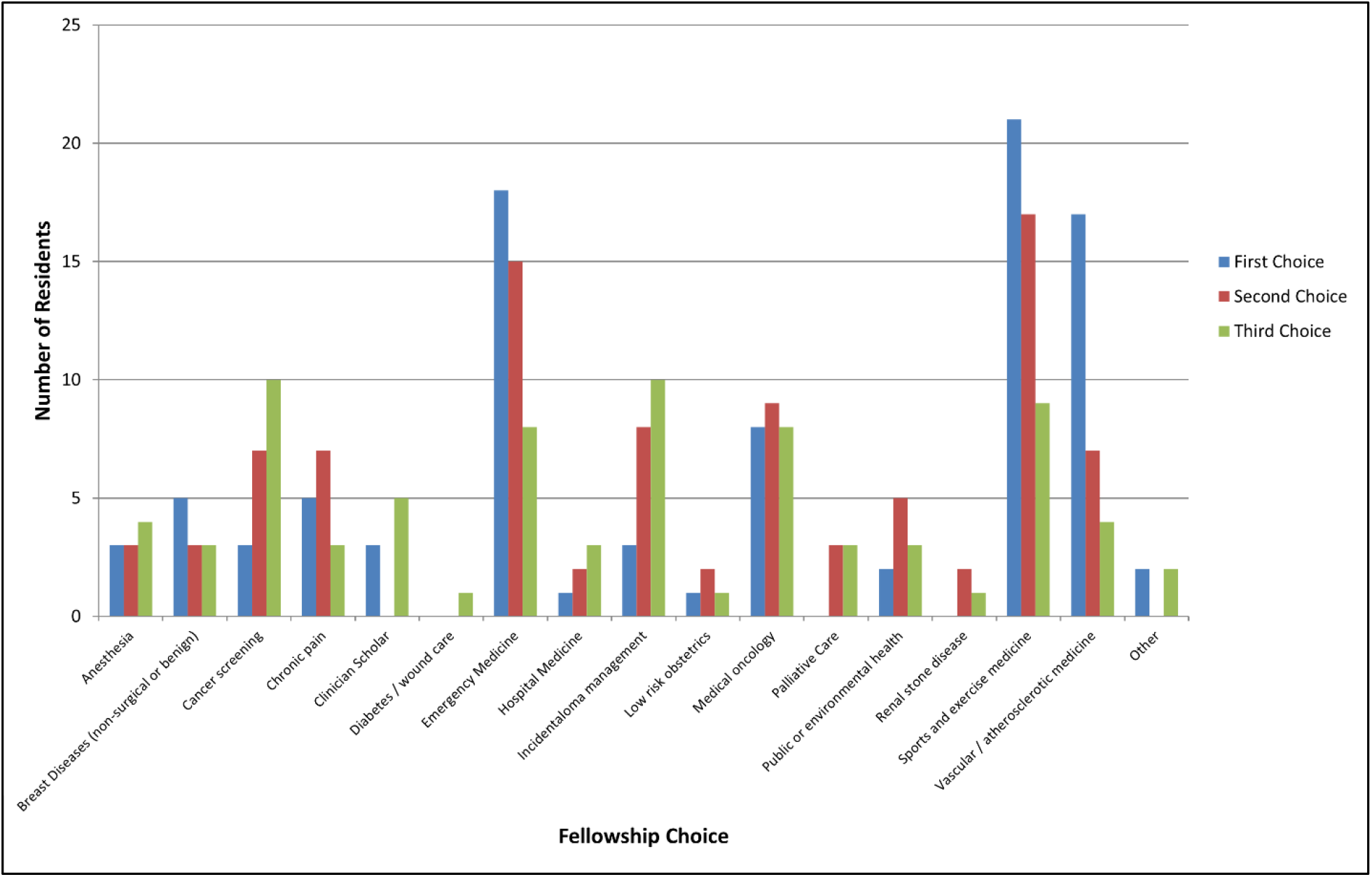
Choices of Expanded Scope of Practice Fellowships among the respondents

63.5% of trainees thought that it was appropriate for radiologists to offer incidentaloma clinics as a component of their practice and 52.9% of trainees deemed themselves capable of offering such a clinic via evidence in the American College of Radiology guidelines.

81.7% of respondents thought it appropriate for radiologists to have access to clinical fellowships to expand their scope of practice.

In addition to these results, we received the following anonymous comments. Comment numbers 5 and 6 were edited for privacy reasons.

1. Thank you for conducting this survey:)
2. Very interesting concept. I find that as radiology continually attempts to add more value to their work, having this type of training and clinics could become useful in the future. It might be difficult to break from the traditional “I sit here and do my reporting”, but with interventional radiologists more and more doing clinics, it might become more of the norm for radiologists to be performing these types of clinics.
3. One could argue that clinical fellowships and increased patient contact are necessary for the survival of radiology as a profession, given the potential for automation in imaging interpretation.
4. If there were an MSK radiology + sports med track, that could be a very compelling program for me.
5. Great project. I think there is a lot of potential for radiologists to expand their practices to include more clinical interaction with patients and colleagues. While I am only a junior resident, breast/women’s imaging has always been one of my interests for that reason. I love radiology but also enjoy seeing patients. Interestingly, I think there is currently a resident from [redacted] that is now doing a clinical/imaging fellowship with the [redacted] group there, and will be able to perform [redacted] etc. and counsel patients upon completion of [redacted] training. More programs like this in the future would be awesome!
6. This is an interesting idea, but I don’t understand (just my lack of knowledge) how this would not put us in direct competition with some of our referring physicians - [redacted]… I think we (as Radiologists) should be looking to add greater value to health care and work closer with our clinical colleagues rather than try to compete with them (and they may compete with us by outsourcing scan reading)… Just my lack of knowledge of the purpose of the survey. But I really appreciate the “out of the box” thinking and maybe this kind of thing would really take off!
7. Interventional radiologists are becoming clinicians in the US and already are clinicians in other parts of the world. My interest in IR leads to a bias regarding my interest in further clinical studies as I am aware that I will already engage in a clinical role in the future.
8. I’m not sure why radiologists want to pursue clinical fellowships after radiology given how weak most radiologists clinical skills are. Clinical medicine changes incredibly rapidly much like radiology and much like radiology takes years to build the clinical intuition and skills to safely practice. Without an integrated continuous component in residency education, I think it will be laughable and more importantly dangerous for radiologists to provide independent clinical care without a lot of experience, support, or well researched guideline support. I think if we want to expand our scope we should decrease general residency training to 3 years (including internship), do 2 years of fellowship training (much like internal medicine), and then spend time on clinical services in that 2 years and then maybe after can do clinical fellowships. Radiology deals with a lot of interventional pain medicine and pursuing clinical fellowships in that or probably palliative type stuff is best. In addition, the MSK people are probably well equipped to undertake additional clinical sports medicine training. Other options like ER or Anesthesia underestimates that amount of work and skill required to practice in those areas (much like people underestimate what it takes to practice radiology well and safe). My $0.02.
9. I have thought about this actually. good initiative and Im sure this is where radiology is heading
10. I feel this is an excellent initiative and something I would absolutely pursue if this was offered. Radiologists have a broad understanding of disease from their training, and I think this especially lends itself to benefits in broad specialties like ER and hospitalist medicine. I can potentially imagine a system where ER-Radiologist physicians are eventually able to sign off on x-ray and ultrasound reads that they interpret live while in the ER or urgent care clinic. This could help improve efficiency and potentially save money by implementing specific fee codes at a subsidized amount for those physicians with capability to bill for the patient encounter and image interpretation.
11. Not an area of personal interest. I chose radiology because I enjoy working/consulting with other medical professionals consulting I did clinical interactions.
12. This is a wonderful idea. Diagnostic Radiology residency provides a comprehensive fund of generalist knowledge and I have often wondered why we are not able to apply this in a clinical setting. Organizations such as the RSNA have been encouraging radiologists to be more visible to patients and clinicians alike, in order to demonstrate our value. Not only would clinical fellowships/practice enable radiologists to demonstrate our value, it would certain add to it. Thanks for doing this!
13. I think including a component of hospitalist and vascular surgery in an Interventional Radiology setting would be appropriate, adding 1 year to fellowship. I also see the value of a pain clinic, as radiologists can provide US and fluoro based procedures. But I’m not really sure diagnostic radiologists have any business in the Emerge or Cancer clinic. If someone wants to do clinical medicine, they should have done a clinical residency.
14. I think ideally mini clinical fellowships (3 months) after radiology fellowships would attract more interest.
15. I would definitely want to do a clinical ER/trauma fellowship.
16. These fellowships should be infolded to any other radiology fellowship, and mandatory at pioneering centres.

## Discussion

We argue that radiologists are qualified to receive referrals from primary care providers for the assessment of lesions that their training enables them to manage including lung nodules, thyroid nodules, small solid renal lesions, cystic renal, hepatic and pancreatic lesions that require monitoring, incidentally detected atherosclerosis or symptomatic atherosclerosis, back pain, amongst many other common clinical problems. The notion of clinical fellowships to expand scope of practice is hence intriguing and well worth the careful consideration of radiologists in North America. In 2017, an imaging residency provides exposure to a range of pathologies rivaled by few other specialties while simultaneously providing training in a wide array of procedures, from simple joint aspirations and injections to line placement and aortic stents.

For example, when a woman of reproductive age presents to the emergency room with pelvic pain, a radiologist who has completed obstetrician-gynecologist rotations in internship, prescribed for those problems, and has performed and interpreted hundreds of pelvic ultrasounds and MRIs has very specific ideas regarding differential diagnosis and workup, pertinent negatives, and lab tests. The knowledge base is competitive with that of a general practitioner and hence the transition of a senior radiology resident or radiology fellow into a clinic environment can be seamless. As radiologists are very much aware, many of these scopes of practice are already defined by nurse practitioners, particularly in community practice, and frequently with a radiologist’s help to avoid complications. In some instances, however, further fellowship training would be helpful to ensure the success and expertise in the expanded scope of practice.

While a remarkable 69% of respondents are interested in seeing patients in clinic as part of their practice, there was also remarkable enthusiasm in taking on a clinical niche via an extra fellowship year including seeing patients in clinic approximately one day a week (i.e. 23.8% of the work week, standard deviation 14.8 hours). We found that 31.7% of respondents had >70% interest in pursuing a clinical fellowship and 13.5% of respondents were completely confident (100%) that they would pursue a clinical fellowship if offered. A majority of residents (~63.5%) felt that it is appropriate for radiologists to undertake incidentaloma clinics without any additional training and more than half (52.9%) felt comfortable offering such a clinic without additional training via support of current American College of Radiology guidelines [9]. Importantly, an overwhelming majority of residents (81.7%) felt that clinical fellowships, as they are defined in the survey, should be accessible to radiologists.

The most popular choices of clinical fellowship in the group were emergency medicine (19.1%), vascular medicine (18.1%) and sports medicine (22.3%). The first two of these choices may be explained partly by the exposure of radiology trainees to the clinical problems in emergency and vascular medicine respectively. Almost one quarter (22.7%) of respondents indicated an interest in interventional radiology fellowships, and we acknowledge that this may have biased our results in favor of clinical fellowships in vascular medicine to some degree, though an almost equal number of residents (23.6%) were unsure of fellowship choice. It is also possible that residents interested in interventional work may have been more likely to respond to the survey.

In addition to the numerical results, we received a number of helpful comments to the survey. These comments are anonymous and we have listed them above in hope that they can generate further discourse. The few negative comments that were received are most helpful and highlight important points. In particular, comment 8 discusses the manner in which expanded scope of practice fellowships might fit into those residency models with shorter general training (i.e. 3 years) and subsequent longer sub-specialty training. The resident also commented that the proposed fellowships echo misconceptions amongst clinicians that radiology can be learned in a year, or via a course, and practiced to a high standard. We respond to these comments only to state that there is strong precedent for one-year fellowships to significantly expand scope of practice in Canada, and that while no training program will ever be perfect, physicians are ultimately personally accountable for their continuing medical education. Cross-specialization must, in our opinion, be a two way street.

In summary, we argue that in 2018, an imaging residency can and should be considered a gateway to any number of clinical specialties. It is appropriate to consider expanded scope of practice fellowships as a means to augment radiology resident’s training. Such programs are likely to be met with considerable enthusiasm.

